# Manufacture of *Clostridioides difficile* spores for experimental infection of human volunteers

**DOI:** 10.1101/2025.09.16.676578

**Authors:** Annefleur D.O. Hensen, Céline Harmanus, Christine Voorvelt, Anoecim R. Geelen, Maryam El Hamdioui, Ed J. Kuijper, Laura Pattacini, Pim Schipper, Inge M. van Amerongen-Westra, Pauline Meij, M.Y Eileen C. Van der Stoep-Yap, Meta Roestenberg, Wiep Klaas Smits

## Abstract

**Background:** Infections with *C. difficile* remain a significant global healthcare problem for which novel (preventive) treatment strategies are urgently needed. Controlled human infection models (CHIMs) can contribute to a better understanding of the disease and accelerate (novel) medicinal product development.

**Methods:** A robust production process for *C. difficile* spores for experimental human administration, was implemented. Direct-release capsules containing viable *C. difficile* spores were formulated. The manufacturing process aligns with good manufacturing practices (GMP) guidelines, including quality controls and release by a qualified person (QP).

**Results:** Selected non-toxigenic and toxigenic *C. difficile* strains were used to create a master cell bank and working cell bank from which multiple batches of purified active substance were produced. Substance was then used to manufacture the final active product. The active product and its intermediate products passed all quality controls for identity, potency, purity and safety, and were individually released by the QP. Stability of active substance and product is confirmed up to 12 months.

**Conclusion:** Although there are no clear standardized international guidelines for challenge material production, we demonstrate that it is feasible to produce *C. difficile* human challenge material for small scale application in CHIM trials. This application of GMP principles to an unconventional production process of a bacterial spore-forming anaerobic challenge agent is an example for the production of challenge material as described in the auxiliary medicinal product guidelines of EMA and FDA.

## Background

*Clostridioides difficile* is the leading cause of healthcare-associated diarrhoea, causing substantial morbidity and mortality. Globally it is responsible for an estimated 15,600 deaths and 284,000 disability adjusted life years (DALYs), representing the primary cause of diarrhoea-related mortality in high-income countries (1). Concerningly, the incidence of community-acquired *C. difficile* infections (CDI) (defined as disease in patients not hospitalized during the 12 weeks prior to diagnosis) is on the rise, almost doubling the past years (2). As such CDI causes a substantial clinical, social and economic burden (3).

An acute episode of CDI is treated with antimicrobial therapy, but alternative therapies are urgently needed because of high relapse rates (approximately 20% after treatment of an initial episode and up to 60% after multiple recurrences) and a rise in antimicrobial resistance (4). Microbial restoration therapies, such as faecal microbiota transplantation (FMT) seem to deliver on the promise of high level sustained prevention of recurrent CDI (5-7). However, the complex interactions between *C. difficile* and the human microbiome remains largely poorly understood, hampering the translation of FMT into a medicinal product. Additionally, animal models cannot fully replicate the human microbiome and its interactions, making preclinical down selection and optimization for novel medicinal products complex. Innovations to accelerate such processes are thus needed.

Controlled human infection models (CHIMs), in which human volunteers are deliberately inoculated with an infectious organism (so-called “challenge agent”) under controlled conditions, are a valuable tool to rapidly evaluate medicinal products (e.g. vaccines) for efficacy in early phase clinical development (8, 9). CHIMs have been developed for a wide variety of pathogens (including parasites, viruses and bacteria) over the past 200 years and CHIM experiments have even led to the registration of a novel cholera vaccine (10). The development of a *C. difficile* CHIM, in which healthy adult volunteers are experimentally exposed to *C. difficile* spores in a controlled clinical environment, would provide the opportunity to identify microbiota and immunological targets for novel (preventive) medicinal products, and ultimately test the preliminary efficacy of such products (e.g. vaccines or microbial restoration therapies).

The development of a *C. difficile* CHIM starts with the production of *C. difficile* challenge material. To ensure safety of CHIM participants a challenge agent which complies with regulatory requirements for human use must be available. Within the EU, challenge agents are classified as auxiliary medicinal products (AxMPs) by the European Medicines Agency (EMA), defined as a medicinal product used for the needs of a clinical trial as described in the protocol, and should be manufactured according to Good Manufacturing Practice (GMP) equivalent standards. GMP guidelines describe the minimum requirements which medicinal products must meet in their manufacturing process to ensure that the medicinal product is 1) of consistent high quality 2) appropriate for their intended use and meets the requirements of the marketing or clinical trial authorization (11).

Aligning the manufacturing process of *C. difficile* challenge material with GMP guidelines, presents several key challenges, particularly in meeting the quality attributes of identity, purity, potency and safety, as outlined in the technical white paper on the quality considerations for challenge agent production for use in CHIM studies (9). Here, we addressed these challenges and describe a robust manufacturing process for challenge material derived from cultures of *C. difficile*, performed in an academic laboratory by trained personnel with an appropriate quality control strategy. The described process for producing encapsulated viable *C. difficile* spores serves as an example of the production of auxiliary medicinal products under the European Medicines Agency (EMA) or U.S. Food and Drug Administration (FDA) regulations.

## Methods

### Manufacturing a *C. difficile* challenge product

Production was performed at a dedicated laboratory for *C. difficile* production of the Experimental Bacteriology research group of the Leiden University Center for Infectious Diseases in the Leiden University Medical Center (LUMC) under the quality management system (QMS) of LUMC’s department of Clinical Pharmacy and Toxicology (KFT) and Center for Cell and Gene Therapy (CCG). The production process of *C. difficile* spores has been established using known and established methods and protocols at the LUMC (12).

The manufacturing process was developed to be adaptable to diverse *C. difficile* strains, including production of non-toxigenic as well as toxigenic *C. difficile* strains. Here, the process to produce challenge material for the non-toxigenic *C. difficile* strain L-NTCD03 (PCR ribotype 416, ST39, clade 4, complete genome GenBank NZ_OX637968.1) and the toxigenic *C. difficile* strain L-TCD-01 (PCR ribotype 020, ST2, clade 1, complete genome GenBank NZ_OY731254.1) is described. Spores were produced in a dedicated anaerobic isolator cabinet (Whitley A135GMP, Don Whitley Scientific, Bingley, UK) in a restricted-access biosafety level 2 laboratory, using food-grade culture media (Soy Peptone A3X (SA3X), Organotechnie, La Courneuve, France) in batch fermentations in sterile single-use polyethylene terephthalate glycol-modified (PETG)-flasks (Sterile Disposable PETG Flask 2000 ml, Thermo Scientific, Rochester, US). Food-grade SA3X was selected as the production medium based on its absence of animal-derived components and its demonstrated capacity for robust growth and high spore yields (data not shown). Spores were purified using aerobic washing with phosphate-buffered saline (PBS) (Fresenius Kabi, Graz, Austria) supplemented with 0.1% Tween 80 (Polysorbatum 80, Duchefa, Haarlem, The Netherlands). Dose-adjusted aliquots (10^4^ and 10^7^ colony forming units (CFU) *C. difficile spores* in buffered 70/30% glycerol (Duchefa, Haarlem, The Netherlands) and PBS solution) were filled into liquid-filled direct-release capsules (Lonza Vcaps® Plus Capsules size 0, Lonza, Colmar, France) using an autoclavable capsule filling system (ProFiller1100, Torpac, Heerlen, The Netherlands). A detailed risk assessment of the *C. difficile* manufacture process and impurities has been conducted. All raw materials used in the manufacturing process are released for their use by a Quality Control (QC) officer. The qualification of all compendial and non-compendial reagents are listed in **Supplementary Table 1**. Non-compendial reagents are qualified based on vendor Certificate of Analysis (CoA) and meet pre-defined acceptance criteria (**Supplementary Table 2 and 3**). The production process was formalized in Standard Operating Procedures (SOPs), authorized by QC, QP and Head of Production.

Quality and in-process controls (QC and IPCs) are described further below and are based on well-established protocols of the Dutch National Expertise Center for *C. difficile* infections (13-15) as well as guidance documents of the European Centers for Disease Control and Prevention (12) and the European Society for Clinical Microbiology and Infectious Diseases (16, 17). All IPCs and QCs were validated or qualified for their intended use. Where possible, IPCs and QCs were performed according to GMP requirements or, if not possible, under ISO15189 standards at the clinical microbiology laboratory of the LUMC. If during culture steps a QC analysis was required, this was performed in a Whitley A35 anaerobic cabinet (Don Whitley Scientific). All QC test results were independently reviewed by a QC officer, and thereafter the master cell bank, working cell bank, active substance and active product were released by a Qualified Person (QP). To verify the robustness of the manufacturing process ≥3 preclinical test runs were performed per challenge strain.

A product dossier was created according to EMA/CHMP/BWP/534898/2008 Rev. 2. for pharmacological products. All raw materials, disposables and excipients were quarantined until release was done by a QC officer.

The product was characterized as an auxiliary medicinal product (regulation EU No 536/2014 (CTR)). The principles of GMP as outlined in Regulation (EU) 536/2014 along with the white paper “Considerations on the Principles of Development and Manufacturing Qualities of Challenge Agents for Use in Human Infection Models” (9) were applied to the production process wherever possible and audited as such.

## Results

### Master cell bank (MCB) and working cell bank (WCB)

The *C. difficile* challenge strains were selected based on considerations previously published (18). The NTCD strain was isolated from faecal material of an asymptomatic nursing-home resident and the TCD strain from faecal material of a symptomatic patient with uncomplicated CDI disease. The selected isolated challenge strains were used to form a master cell bank (MCB), the starting material for the production of the active product (**Figure 1**). MCB has been prepared under good laboratory practice (GLP) conditions according to the SOPs of the Dutch National Expertise Center for *C. difficile* infections, which is an ISO-15189 certified laboratory. In short, the MCB was created through enrichment culture in CDEB MOD (*C. difficile* enrichment modified broth, Mediaproducts BV, Groningen, The Netherlands), inoculated onto CLO plates (*Clostridioides difficile* selective agar plates, bioMérieux, Marcy-l’Étoile, France), followed by re-culturing on a tryptic soy sheep blood (TSS) plate (bioMérieux, Marcy-l’Étoile, France), after which a colony was transferred to glycerol for storage at -80°C.

**Figure 1.**
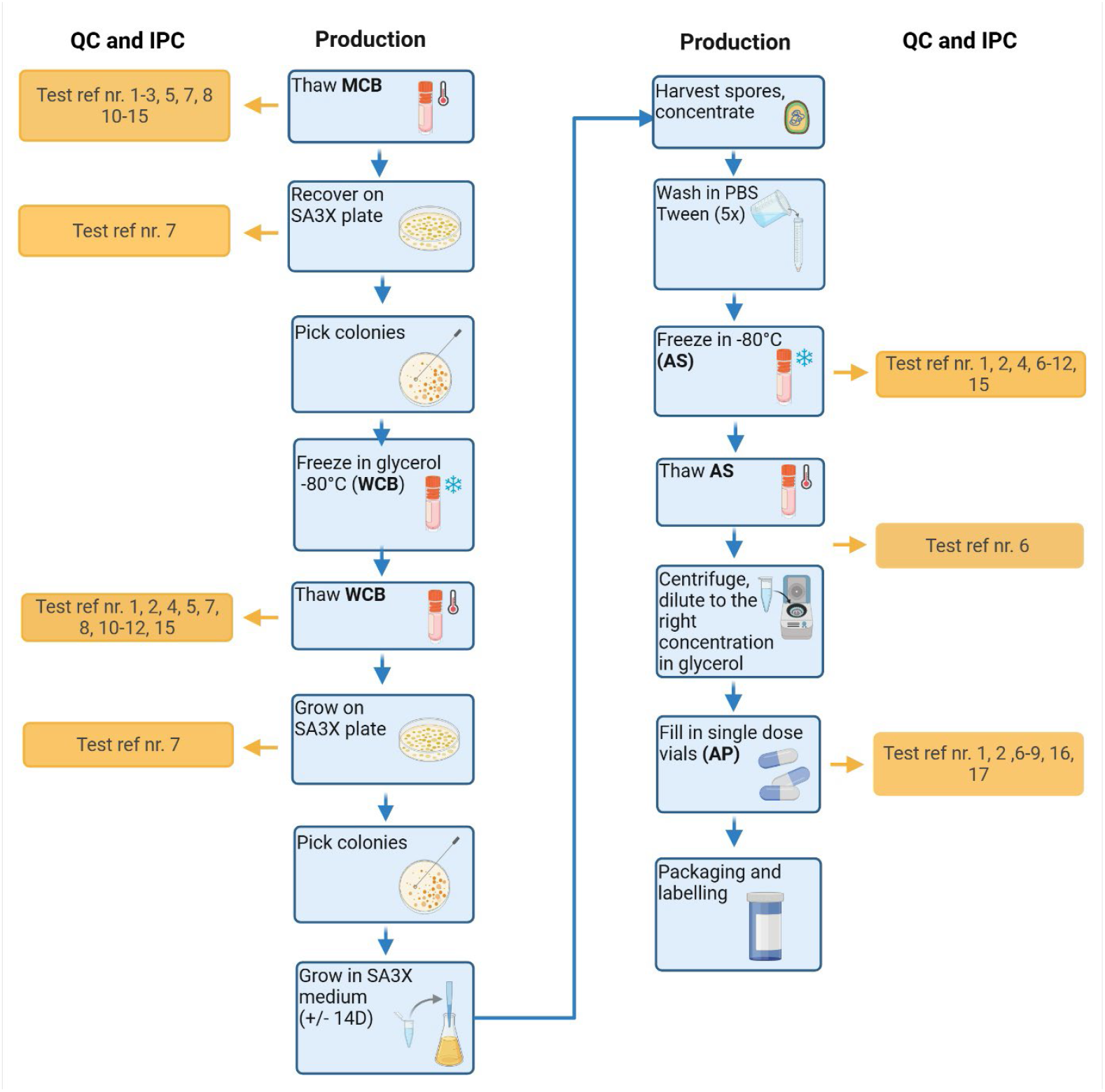
Flow chart of the *C. difficile* challenge material production process. For the test reference numbers please see **Table 1**. Abbreviations: QC= quality control, IPC: in-process control, MCB = master cell bank, SA3X= Soy Peptone A3X; WCB= working bank, PBS = phosphate buffered saline, AS = active substance, AP= active product. Figure created with BioRender.com.

To produce the working cell bank (WCB) intermediate product (**Figure 1**), MCB was thawed and plated out on a SA3X plate for expansion of *C. difficile*. After incubation of the plate, colonies were picked and subsequently the WCB was stocked in a glycerol/SA3X solution and stored in a -80°C freezer.

The MCB and WCB were released for further processing after undergoing successful quality control testing for confirmation of identity, potency, purity and safety of the products and passing all acceptance criteria (**Table 1**). NGS testing additionally showed stability of the challenge strain genome, with no evidence of laboratory-driven adaptation of the original strain (data not shown).

**Table 1.** In-process control and quality control tests, methods and acceptance criteria applied to the master cell bank (MCB), working cell bank (WCB), active substance (AS) and active product (AP). Abbreviations: PCR= polymerase chain reaction; Y= yes; N=no; RT= ribotype; GLP= good laboratory practice; NGS= next generation sequencing; ST= sequence type; CFU= colony forming unit; *gluD*= glutamase dehydrogenase; TSS=Tryptic Soy Sheep blood; NTCD= non-toxigenic *C. difficile*, TCD= toxigenic *C. difficile*, neg= negative, pos=positive, MZ= metronidazole; VA= vancomycin; FDX= fidaxomicin. Figure created with BioRender.com.

| Test ref. nr | IPC/QC | Description | Method | Acceptance criteria | Applied to |  |  |  |
| --- | --- | --- | --- | --- | --- | --- | --- | --- |
| Identity |  |  |  |  | MCB | WCB | AS | AP |
| 1 | QC | Species PCR* | *gluD* PCR | Positive | Y | Y | Y | Y |
| 2 | QC | Ribotyping* | Capillary ribotyping | RT identical to the GLP clinical isolate | Y | Y | Y | Y |
| 3 | QC | Sequencing | NGS (complete genome/long read) | Species: *Clostridioides difficile*<br>ST: identical to the GLP clinical isolate | Y | N | N | N |
| 4 | QC | Sequencing | NGS (short read/Illumina) | Species: *Clostridioides difficile*<br>ST: identical to the GLP clinical isolate | N | Y | Y | N |
| Potency |  |  |  |  | MCB | WCB | AS | AP |
| 5 | QC | Cell count | CFU count | >100 CFU/ml | Y | Y | N | N |
| 6 | QC | Spore count* | Spore enumeration | S: > 1.6 x 10<sup>6</sup> spores/mL<br>P: targeted dose 10<sup>4</sup> (range 10<sup>3</sup>-10<sup>5</sup>)<br>P targeted dose 10<sup>7</sup> (range 10<sup>6</sup>-10<sup>8</sup>) | N | N | Y | Y |
| Purity |  |  |  |  | MCB | WCB | AS | AP |
| 7 | IPC | Visual appearance of colonies* | Visual inspection of colonies on TSS plate | Visual appearance consistent with *C. difficile* species | Y | Y | Y | Y |
| 8 | QC | Contamination with aerobic bacteria* | Aerobic culturing on a TSS plate | No growth of aerobic bacteria (<10 CFU/mL) | Y<sup>1</sup> | Y | Y | Y |
| 9 | QC | Measure protein and DNA content* | Colorimetric assay | Protein: < 10 ug/uL<br>dsDNA: < 10 ng/uL | N | N | Y | Y |
| Safety |  |  |  |  | MCB | WCB | AS | AP |
| 10<sup>2</sup> | QC | Presence of toxin | Multiplex qPCR for toxin A, B and binary toxin | *cdtA*, *cdtB*: NTCD and TCD neg<br>*tcdA*, *tcdB*: NTCD neg, TCD pos<br>*gluD*, 16S rRNA: NTCD and TCD pos | Y | Y | Y | N |
| 11 | QC | Antimicrobial susceptibility testing (MZ en VA) | E-test | MZ: <0.5 mg/L<br>VA: <0.5mg/L | Y | Y | Y | N |
| 12 | QC | Antimicrobial susceptibility testing (FDX) | Agar dilution | <0.5 mg/L | Y | Y | Y | N |
| 13 | QC | Microbial contaminants | Presence of pathogenic bacterial contaminants | PCR negative for *Salmonella*, *Shigella*, *Campylobacter*, *Yersinia*, *Aeromonas* and *Plesiomonas shigelloides* spp. | Y | N | N | N |
| 14 | QC | Microbial contaminants | Presence of pathogenic viral contaminants | PCR negative for norovirus, astrovirus, sapovirus, adenovirus and rotavirus | Y | N | N | N |
| 15 | QC | NGS for adventitious agents | NGS for DNA | Positive assignment as *Clostridioides difficile*; no discovery with >0.3% of mapped reads to pathogenic species | Y | Y | Y | N |
| Capsule controls |  |  |  |  | MCB | WCB | AS | AP |
| 16 | QC | Capsule integrity* | Visual inspection | Capsules are properly locked<br>Capsules are whole and not dented | N | N | N | Y |
| 17 | QC | Fill weight and volume* | Weight | Volume: 500 uL<br>Fill weight: 0.70 gram (+/-0.02 gram) | N | N | N | Y |

<sup>1</sup> Only performed on the TCD MB. <sup>2</sup> For test reference number 10, the acceptance criteria listed are based on L-NTCD03 and L-TCD-01 (see Methods) and may differ if other strains are used.
\* These tests were also performed for stability testing of the active product.

### Active substance (AS)

To produce purified spores in solution (**Figure 1**), which constitute the active substance (AS), the WCB was recovered on a SA3X agar plate and re-streaked on a fresh SA3X plate to increase cell yield. Subsequently, a starter culture was generated in liquid SA3X medium under anaerobic conditions using the anaerobic isolator cabinet. The optical density (OD) of the cultures was then measured and diluted in fresh SA3X medium to a starting OD of 0.05. The diluted cultures were incubated in shaking Erlenmeyer flasks for 14 days to produce spores. In the production medium, spore yield increases during the first week (from 0 CFU/mL to >10^6^/mL); a 14-day time frame was selected to ensure maximum reproducibility of spore yield. We found no significant difference in spore yield between different culture volumes up to a liter-scale, demonstrating robustness of the method (data not shown). After 14 days, cells were pelleted by centrifugation and washed 5 times with PBS/0.1% Tween 80 to remove residual vegetative cell debris. All batches of the active substance yielded a high spore count (**Figure 2**), meeting the acceptance criterion for spore count (>1.6×10^6^ spores/mL). Active substance was stored at -80°C.

**Figure 2.**
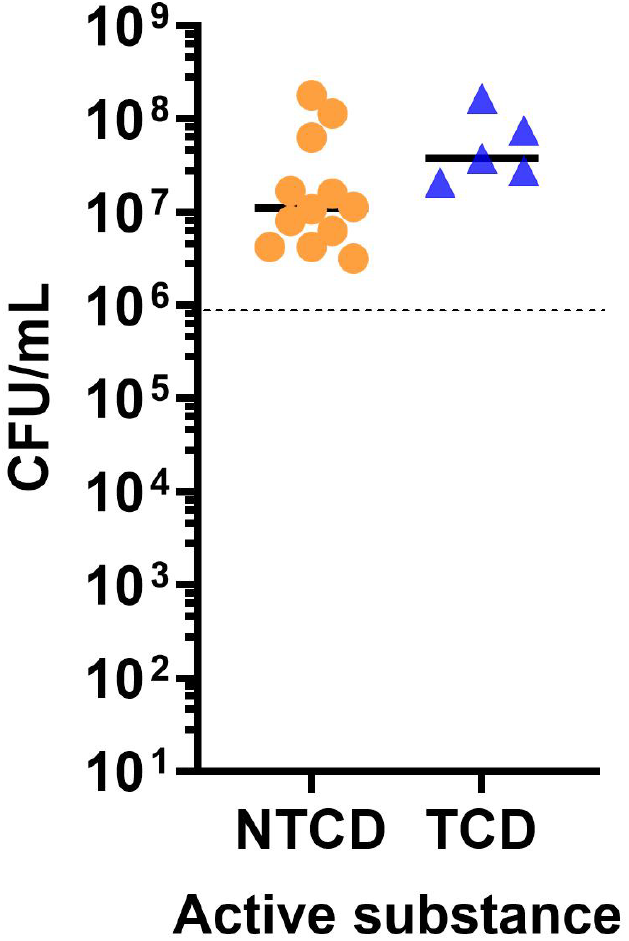
Spore yield (CFU/mL) of different batches of active substance (NTCD and TCD). The dotted line indicates the defined acceptance criterion. The NTCD strain used is L-NTCD03 and the TCD strain used is L-TCD-01. Abbreviations: NTCD= non-toxigenic *C. difficile*, TCD= toxigenic *C. difficile*, CFU/mL= colony forming units per milliliter.

The active substance was released for further processing after undergoing successful quality control testing and passing all acceptance criteria as indicated in **Table 1**.

### Active Product (AP)

The active product consists of a 500-μl suspension containing 10^4^ or 10^7^ CFU *C. difficile* spores formulated in buffered 70/30% (v/v) glycerol and PBS encapsulated in liquid-filled direct-release capsules for oral administration. The dosing of 10^4^ and 10^7^ CFU spores were selected based on previously published experimental NTCD studies (19, 20).

To produce the active product, active substance was thawed and adjusted to the right concentration in formulation buffer after which 500-μl volumes were dispensed in liquid-filled direct-release Lonza Vcaps®. Capsules were stored at room temperature (15-25°C) and quarantined until release.

The active product was checked for identity, potency, purity and safety, and released after passing all acceptance criteria (**Table 1**).

### Stability of Active Product

Stability testing was performed on the active substance and active product at multiple timepoints according to a predefined stability plan (1, 3, 6, 12 and 24 months). Stability testing consists of: 1) identity testing by *gluD* PCR and ribotyping PCR 2) potency testing by spore enumeration (**Figure 3**) 3) purity testing by visual inspection of colonies on a TSS plate, colorimetric measurement of protein and DNA content and aerobic plating and capsule controls by visual inspection of capsule integrity and measuring fill weight and volume (**Table 1**).

**Figure 3.**
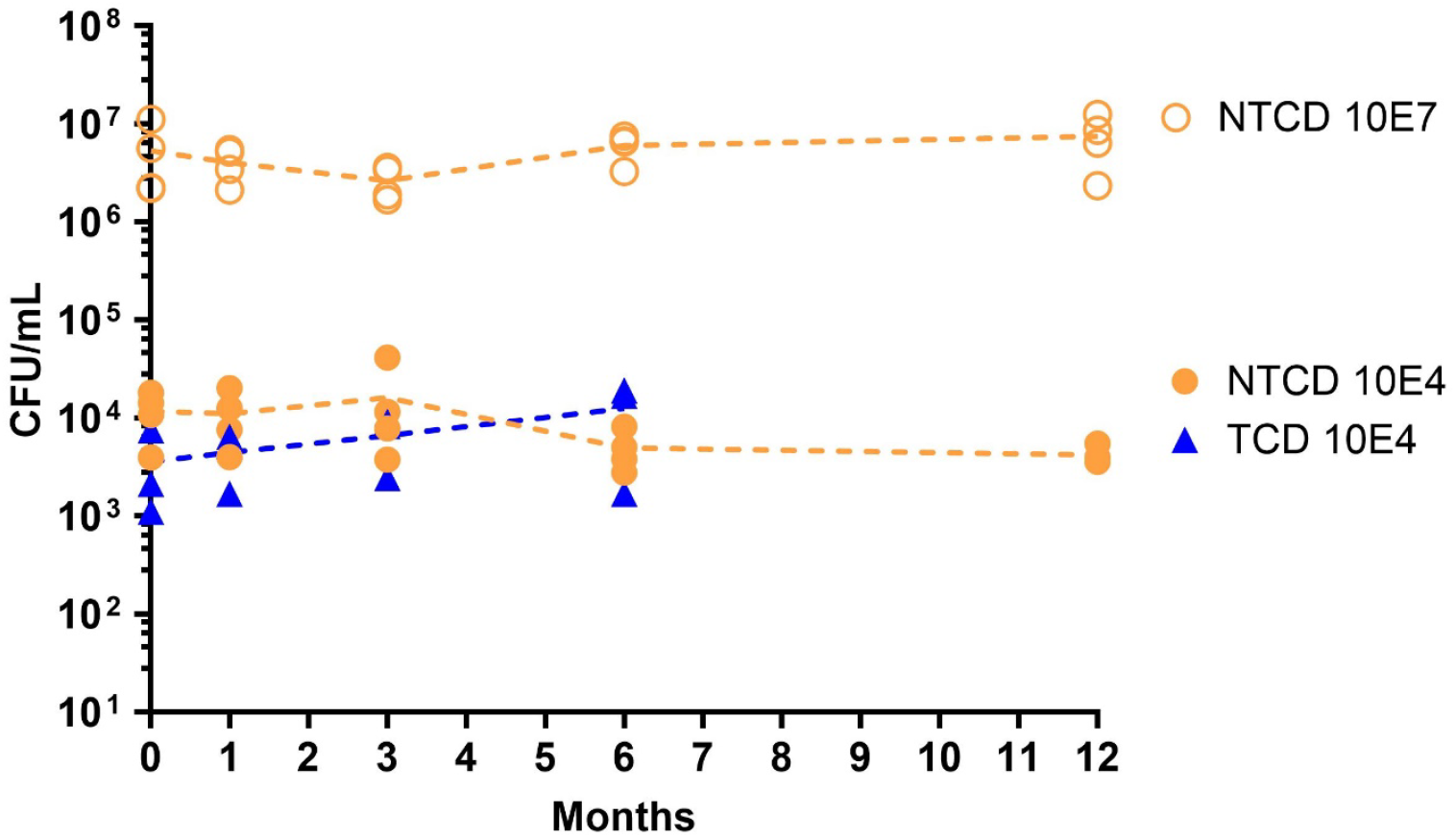
Spore yield stability of multiple batches of active product (capsules) over time (NTCD and TCD). The NTCD strain used is L-NTCD03 and the TCD strain used is L-TCD-01.TCD stability testing is ongoing. Abbreviations: NTCD= non-toxigenic *C. difficile*, TCD= toxigenic *C. difficile*, CFU/mL= colony forming unit per milliliter.

We observed stability, with limited or no decrease in plating efficiency over a period of 12 months so far for the NTCD active product, and up to 6 months for the TCD active product (**Figure 3**). All other stability tests also met the predefined criteria (**Table 1**) at these timepoints. Stability of the active substance aligns with the stability observed of the active product. The observed stability at multiple timepoints across different production batches confirms robustness of stability across the batches. The expiry date of the (N)TCD active product is adjusted to the stability data. Notably, stability of the TCD product to date strongly resembles the NTCD product, suggesting a similar stability profile. Based on these observations, we expect that the actual shelf life of the challenge product extends well beyond the current stability data. Together this demonstrates that the active product has a sufficiently long shelf life (stability) to enable its use in challenge studies following production.

### In-process controls and quality controls

The in-process controls (IPCs) and quality controls (QCs) performed to ensure identity, potency, purity and safety during the production process of *C. difficile* challenge material are scheduled in **Figure 1** and **Table 1**. A short description of the tests is given below.

#### Identity

##### Species PCR

Confirmation of the presence of cells/spores belonging to the species *C. difficile* was obtained by performing a *gluD* PCR on isolated total DNA. The *gluD* PCR contains highly specific primers which allow amplification of a 158-bp fragment of glutamate dehydrogenase gene (*gluD* gene), after which the fragment can be detected using a specific *gluD* probe in a real time PCR assay (12).

##### Ribotyping

To identify and confirm the PCR ribotype of the *C. difficile* challenge material capillary ribotyping PCR was performed, in which the 16S-23S intergenic spacer regions on the *C. difficile* chromosome are amplified by PCR and visualized on a capillary gel electrophoresis system (14). The resulting banding pattern is matched to a library of reference patterns, using BioNumerics 7.6 software (bioMérieux, Marcy-l’Étoile, France).

##### Complete genome sequencing

A reference sequence for the challenge strains was generated using long-read circular consensus sequencing (Pacific Bioscience Sequel II) and automated assembly using established protocols (21, 22). Reference assembly of the 150-bp paired-end Illumina reads of this sequence revealed no single nucleotide polymorphisms. Analysis of the complete genome sequence allowed for assessment of genome characteristics such as the presence or absence of specific toxin and antimicrobial resistance genes.

##### Short read sequencing

Short-read next generation sequencing (Illumina Miniseq, NextSeq 500 or Novaseq Xplus) was additionally performed on the working bank and active substance; the results of a core genome analysis using Ridom SeqSphere+ (Ridom GmbH, Münster, Germany) confirmed that the clade (clonal complex, CC) and complex type (CT) of the strain matches the reference genome sequence.

#### Potency

##### Total cell count

To enumerate viable bacterial cells in the master and working cell bank, serial dilutions were cultured on BHI (brain heart infusion) plates. The total cell count is based on the geometric mean minus two times standard deviation of log transformed values obtained from the colony forming unit (CFU) enumeration.

##### Total spore count

The quantity of viable spores was assessed by plating out serial dilutions of the product onto BHI plates supplemented with 0.1% of the germinant sodium taurocholate (Thermo Fisher Scientific, Lancashire, UK) and enumerating the CFUs after 48h. The total spore count is based on the geometric mean minus two times standard deviation of log transformed values obtained from the CFU enumeration. Production yield of active substance was targeted at > 1.6 x 10^6^ spores/mL, but generally produced >10^7^ spores/mL. In our studies, the active product was formulated in capsules with a targeted dose of 10^4^ or 10^7^ spores.

#### Purity

##### Visual appearance of colonies

Routinely, *C. difficile* cells or spores were anaerobically cultured on a TSS plate for 72h. Plates were visually inspected and all colonies on the plate should macroscopically be morphologically consistent with *C. difficile*, which required confirmation by a second person.

##### Contamination with aerobic bacteria

To confirm absence (<10 CFU/mL) of aerobic bacterial contaminants, routine aerobic culture was performed on non-selective TSS agar plates.

##### Measuring protein and DNA content

To quantify remaining vegetative *C. difficile* cell remnants, e.g. free protein or DNA, in the active substance and product, colorimetric measurements (Qubit dsDNA HS Assay and Qubit Protein Assay, LifeTechnologies, Eugene, Oregon, US) were performed. The limits set at ∼ 10-fold the minimum value observed after repeated washing (please refer to **Table 1**).

#### Safety

##### Presence of toxin gene

A multiplex PCR for genes encoding toxin A (*tcdA*), toxin B (*tcdB*) and binary toxin (*cdtA* and *cdtB*) genes was performed on total DNA extracted from the samples. As a DNA isolation control, the 16S rRNA and *gluD* gene were also tested, which should show a positive result.

##### Antimicrobial susceptibility

To ensure susceptibility to antibiotics which may be used for rescue treatment (fidaxomicin, vancomycin and metronidazole), antimicrobial susceptibility tests are performed. Susceptibility for fidaxomicin (Selleckchem, Köln, Germany) was tested using the agar dilution method (23) whereas for vancomycin and metronidazole E-tests (bioMérieux, Marcy-l’Étoile, France) were used. Breakpoints are defined based on EUCAST criteria (24).

##### Microbial contaminants of the master cell bank

To test for contaminants with gastrointestinal pathogens, total DNA from the master cell bank was tested by qPCR for the presence of gastrointestinal pathogens, i.e. the bacterial pathogens *Salmonella, Shigella, Campylobacter, Yersinia, Aeromonas* and *Plesiomonas shigelloides spp*. and the viral pathogens adenovirus, norovirus, rotavirus, sapovirus and astrovirus.

##### Next-generation sequencing (NGS) for bacterial adventitious agents

An additional metagenomic analysis was performed to identify possible contaminants in a non-targeted manner. Short-reads from the Illumina sequencing (see above) were analysed through the GenomeDetective tool (EmWeb bv, Herent, Belgium). This analysis classifies reads to bacterial species and allows to confirm both identity (assignment: *Clostridioides difficile*) and assess the level of selected bacterial pathogens (25). As metagenomic analysis is prone to result in spurious identifications (for instance because of DNA segments that are shared between bacterial species), acceptance criteria were validated on the basis of analysis of simulated reads derived from reference genome sequence.

#### Capsule controls

##### Capsule integrity

Capsules from the beginning, middle and end of the manufacturing process were inspected for integrity by visually inspecting the capsules: both capsule parts should be aligned, show no deformations and no liquid should be leaking out when tilted.

##### Fill weight and volume

The fill weight of the active product capsules is measured on an analytical balance; each capsule is filled with 500ul which should align with a weight of 700mg +/- 20 gram.

## Conclusion and Discussion

To summarize, we demonstrate feasibility to produce *C. difficile* challenge material for use in CHIM studies at small scale aligning to GMP principles including appropriate quality controls and release. This application of GMP principles to an unconventional production process of a bacterial spore-forming anaerobic challenge agent is a representation for the production of bacterial challenge agents as described in the auxiliary medicinal product guidelines of EMA and FDA.

Challenge material is not classified as a medicinal product and specific international challenge material production guidelines are lacking. The EMA identifies challenge agents as auxiliary medicinal products (AxMPs) and states that when the AxMP is not authorized it shall be manufactured according to GMP *or* at least an equivalent standard (26). In the United Kingdom, the manufacturing of challenge agents is not regulated. In the US the challenge agent is considered a biological product and is subject to regulation under federal law by the FDA. The challenge agent must therefore 1) be manufactured under GMP conditions where possible, 2) satisfy the FDA regulations of safety, purity and potency and 3) have detailed information on the provenance and manufacture as part of the required Investigational New Drug Application (IND).

To ensure the quality attributes of identity and safety NGS was applied for full characterization of the challenge strain. Additionally, applying NGS for the detection of adventitious agents enhances the QC testing of challenge material facilitating the identification of potential other bacterial pathogens in the active product beyond *Ph. Eur*. requirements. To safeguard the level of contamination with adventitious agents, acceptance criteria must be predefined. This can be challenging, as metagenomic analysis is prone to spurious identifications (e.g. due to DNA segments shared between bacterial species). This is of particular concern for *C. difficile* spore challenge material, as DNA isolation from spores is difficult and typically yields low-biomass DNA samples. We established acceptance criteria for detecting adventitious agents through NGS by analysing simulated reads generated from the reference genome using a well-characterized pipeline, resulting in a cut-off of fewer than 0.3% of mapped reads aligning to pathogenic species. Such criteria may require adaptation for other species. In the case of spurious NGS identifications, a quantitative 16S micelle PCR (27) can provide additional value by enabling both the identification and quantification of the bacterial composition.

Secondly, to ensure the quality attribute of safety, food-grade media and GMP-grade raw materials– formulated to avoid the use of animal-derived products where possible, thereby mitigating the risk of potentially important diseases such as transmissible spongiform encephalopathies (TSE) - are required during challenge material production. However, the culture media routinely used for *C. difficile* growth and sporulation contain animal-derived constituents, and conventional spore purification techniques involve reagents that are not indicated for use in a spore production process or for oral ingestion (e.g. non-ionic X-ray contrast agents). Therefore, during the manufacturing process development, the normally used brain heart infusion (BHI) medium was replaced by a soy peptone (SA3X) medium and for spore purification washing with PBS-0.1% Tween was established. Moreover, the dedicated manufacturing area was adapted to include appropriate control measures, equipment validation and a cleaning protocol to prevent cross-contamination. Although GMP guidelines may stipulate production in a controlled environment (e.g. a clean room), the inherent resistance and persistence of *C. difficile* spores present significant challenges for decontaminating such facilities. Notably, the guidance document on the production of challenge agents (9) states that a clean room is not mandatory, but the manufacturing area should be a part of the contamination control strategy and the level of environmental control for particulate and microbial contamination should be tailored to the manufacture of the challenge agent and production step, considering the potential level of contamination of starting materials and the risks for the final batch of the challenge agent. Since *C. difficile* requires anaerobic conditions during production, an anaerobic isolator cabinet was needed as the manufacturing area, and to maintain a robust and reproducible manufacturing process a qualified anaerobe cabinet (GMP grade A) was selected. The isolator underwent full qualification (installation, operational and performance qualification). Thereby this manufacturing area, provided not only the necessary environment for spore production but also ensured environmental control, including regulation of particulate and microbial contamination through controlled, HEPA-filtered, unidirectional airflow.

Thirdly, to ensure the quality attribute of potency, the route of administration, storage and transport conditions of the challenge agent must be carefully considered. Oral capsules were chosen as the route of administration to mimic the natural infection route, while simultaneously maintaining control over the number of spores reaching the gastrointestinal tract. As *C. difficile* spores need exposure to primary bile acids in the duodenum for germination (28, 29), liquid-filled direct-release capsules (instead of enteric capsules) were chosen as they release the spores in the stomach or shortly after. 70/30% (v/v) buffered glycerol/PBS was selected as the formulation buffer for its optimal balance between feasibility to formulate the spore suspension and prevention of downstream capsule leakage. The capsules were preserved at room temperature (15-25 °C) to avoid structural changes in the content and integrity of the capsules which occur upon freezing. Extensive testing of different capsule types demonstrated best performance of Lonza Vcaps® liquid-filled direct-release capsules in terms of secure containment of liquids and semi-solids, stability and preventing leakage at room temperature.

Applying the GMP principles to *C. difficile* spore challenge material for clinical use highlights the need for collaboration and thorough risk analysis among various disciplines, including pharmacists, microbiologists, clinicians and technicians. This collaboration has led to a highly controlled production of *C. difficile* spore challenge material in an academic setting. Building on these efforts, the *C. difficile* challenge material has been approved by the Medical Research Ethics Committee to use for colonisation (with non-toxigenic *C. difficile* spores, ClinicalTrials.gov NCT05693077) and infection (with toxigenic *C. difficile* spores, ClinicalTrials.gov NCT06702345) of healthy volunteers in a proof-of-concept clinical trial to find a safe and infectious dose. Establishing a CHIM for *C. difficile* will be a critical advancement in improving our understanding of *C. difficile* infections as well as accelerating (novel) product development to battle the rising incidence and global burden of this complex disease.

## Contribuon

*Writing-original draft* (lead): AH, *Writing original draft (supporting):* MR and WKS, *Writing-review and editing*: EK, PS, PM, ES, IA, CH, CV, LP *Conceptualization:* WKS, MR, EK, ES, PM and AH, *Investigation*: CH, CV, AG and ME, *Methodology*: ES, PM, MR, WKS, EK, CH, CV, AG and ME, *Resources*: PS, PM, IA, ES, EK and WKS *Project administration*: LP, *Validation*: ES, PM, IA and PS, *Supervision*: WKS and MR.

## Acknowledgements

We gratefully acknowledge the contributions of all partners involved in Work Package 10 (Subtopic 2) of the Inno4Vac project, an Innovative Health Initiative-funded consortium.

## Declaraon of compeng interest

Wiep Klaas Smits performs research which is funded in part by a public-private partnership with Acurx Pharmaceuticals unrelated to the work presented here. All other authors declare no competing interests.

## Supplementary Data

**Supplementary Table 1.** Qualifications of the raw materials used during the production process of *C. difficile* challenge material.

| Material | Manufacturer | Grade | Release criteria | Reference |
| --- | --- | --- | --- | --- |
| Agar | ThermoFisher Scientific | Ultrapure | CoA | Release statement |
| SA3X medium | Organotechnie S.A.S. | Research | CoA | Release statement |
| Tween 80 | Duchefa | Ph. Eur | CoA | Release statement |
| Glycerol | Duchefa | Ph. Eur | CoA | Release statement |
| Phosphate buffered saline (PBS) | Fresenius Kabi | Ph. Eur. | CoA | Release statement |
| Water for injection | Fresenius Kabi | Ph. Eur. | SmPC | Release statement |

**Supplementary Table 2.** Acceptance criteria of Agar medium.

| Test | Acceptance criteria |
| --- | --- |
| Ash | ≤6.5% |
| Gel point (1.5% gel) | 32-38°C |
| Gel strength (1.5% gel) | 550-950 g/cm<sup>2</sup> |
| Gel pH (1.5%) after autoclaving | pH 6.0-7.5 |
| Melting point (1.5%) | 80-90°C |
| Moisture | ≤12% |

**Supplementary Table 3.** Acceptance criteria of SA3X medium.

| Test | Acceptance criteria |
| --- | --- |
| Appearance | Beige powder |
| Solubility in water at 5% | Complete |
| pH (5% solution) | 7.4 |
| Loss on drying | 5% |
| Total nitrogen TN | 9.3% |
| A-Amino nitrogen AN | 3.5% |
| AN/TN x 100 | 38 |
| Residue on ignition | 17% |
| Chloride (NaCl) | 0.4% |
| TAMC | ≤10000/g |

## References

1. Collaborators GBDDD. Global, regional, and national age-sex-specific burden of diarrhoeal diseases, their risk factors, and aetiologies, 1990-2021, for 204 countries and territories: a systematic analysis for the Global Burden of Disease Study 2021. Lancet Infect Dis. 2025;25(5):519–36 doi:10.1016/S1473-3099(24)00691-1

2. ECDC. Clostridioides difficile infections. Annual epidemiological report for 2018−2020. Stockholm: ECDC; 2024.

3. Jones AM, Kuijper EJ, Wilcox MH. Clostridium difficile: a European perspective. J Infect. 2013;66(2):115–28 doi:10.1016/j.jinf.2012.10.019

4. Smits WK, Garey KW, Riley TV, Johnson S. Clostridioides difficile is a bacterial priority pathogen. Anaerobe. 2025;93:102965 doi:10.1016/j.anaerobe.2025.102965

5. van Nood E, Vrieze A, Nieuwdorp M, Fuentes S, Zoetendal EG, de Vos WM, et al. Duodenal infusion of donor feces for recurrent Clostridium difficile. N Engl J Med. 2013;368(5):407–15 doi:10.1056/NEJMoa1205037

6. Feuerstadt P, Louie TJ, Lashner B, Wang EEL, Diao L, Bryant JA, et al. SER-109, an Oral Microbiome Therapy for Recurrent Clostridioides difficile Infection. N Engl J Med. 2022;386(3):220–9 doi:10.1056/NEJMoa2106516

7. Khanna S, Assi M, Lee C, Yoho D, Louie T, Knapple W, et al. Efficacy and Safety of RBX2660 in PUNCH CD3, a Phase III, Randomized, Double-Blind, Placebo-Controlled Trial with a Bayesian Primary Analysis for the Prevention of Recurrent Clostridioides difficile Infection. Drugs. 2022;82(15):1527–38 doi:10.1007/s40265-022-01797-x

8. Roestenberg M, Hoogerwerf MA, Ferreira DM, Mordmuller B, Yazdanbakhsh M. Experimental infection of human volunteers. Lancet Infect Dis. 2018;18(10):e312–e22 doi:10.1016/S1473-3099(18)30177-4

9. La CB, Alan; Trouvin, Jean-Hugues; Depraetere, Hilde; Neels, Pieter; R. Talaat Kawsar; et al. Considerations on the principles of development and manufacturing qualities of challenge agents for use in human infection models. Wellcome Trust. 2022

10. Chen WH, Cohen MB, Kirkpatrick BD, Brady RC, Galloway D, Gurwith M, et al. Single-dose Live Oral Cholera Vaccine CVD 103-HgR Protects Against Human Experimental Infection With Vibrio cholerae O1 El Tor. Clin Infect Dis. 2016;62(11):1329–35 doi:10.1093/cid/ciw145

11. EMA. 2025 [cited 2025 26-06-2025]. Available from: https://www.ema.europa.eu/en/human-regulatory-overview/research-development/compliance-research-development/good-manufacturing-practice.

12. ECDC. Laboratory procedures for diagnosis and typing of human Clostridium difficile infection. Stockholm: ECDC; 2018.

13. Baktash A, Corver J, Harmanus C, Smits WK, Fawley W, Wilcox MH, et al. Comparison of Whole-Genome Sequence-Based Methods and PCR Ribotyping for Subtyping of Clostridioides difficile. J Clin Microbiol. 2022;60(2):e0173721 doi:10.1128/JCM.01737-21

14. Fawley WN, Knetsch CW, MacCannell DR, Harmanus C, Du T, Mulvey MR, et al. Development and validation of an internationally-standardized, high-resolution capillary gel-based electrophoresis PCR-ribotyping protocol for Clostridium difficile. PLoS One. 2015;10(2):e0118150 doi:10.1371/journal.pone.0118150

15. RIVM. Clostridioides (Clostridium) difficile 2024 [cited 2025 24-06-2025]. Available from: https://www.rivm.nl/clostridioides-difficile.

16. van Prehn J, Crobach MJT, Baktash A, Duszenko N, Kuijper EJ. Diagnostic Guidance for C. difficile Infections. Adv Exp Med Biol. 2024;1435:33–56 doi:10.1007/978-3-031-42108-2_3

17. van Prehn J, Reigadas E, Vogelzang EH, Bouza E, Hristea A, Guery B, et al. European Society of Clinical Microbiology and Infectious Diseases: 2021 update on the treatment guidance document for Clostridioides difficile infection in adults. Clin Microbiol Infect. 2021;27 Suppl 2:S1–S21 doi:10.1016/j.cmi.2021.09.038

18. Hensen ADO, Vehreschild M, Gerding DN, Krut O, Chen W, Young VB, et al. How to develop a controlled human infection model for Clostridioides difficile. Clin Microbiol Infect. 2025;31(3):373–9 doi:10.1016/j.cmi.2024.08.025

19. Villano SA, Seiberling M, Tatarowicz W, Monnot-Chase E, Gerding DN. Evaluation of an oral suspension of VP20621, spores of nontoxigenic Clostridium difficile strain M3, in healthy subjects. Antimicrob Agents Chemother. 2012;56(10):5224–9 doi:10.1128/AAC.00913-12

20. Gerding DN, Meyer T, Lee C, Cohen SH, Murthy UK, Poirier A, et al. Administration of spores of nontoxigenic Clostridium difficile strain M3 for prevention of recurrent C. difficile infection: a randomized clinical trial. JAMA. 2015;313(17):1719–27 doi:10.1001/jama.2015.3725

21. Ducarmon QR, van der Bruggen T, Harmanus C, Sanders I, Daenen LGM, Fluit AC, et al. Clostridioides difficile infection with isolates of cryptic clade C-II: a genomic analysis of polymerase chain reaction ribotype 151. Clin Microbiol Infect. 2023;29(4):538 e1–e6 doi:10.1016/j.cmi.2022.12.003

22. Nooij S, Plomp N, Sanders I, Schout L, van der Meulen AE, Terveer EM, et al. Metagenomic global survey and in-depth genomic analyses of Ruminococcus gnavus reveal differences across host lifestyle and health status. Nat Commun. 2025;16(1):1182 doi:10.1038/s41467-025-56449-x

23. Freeman J, Sanders I, Harmanus C, Clark EV, Berry AM, Smits WK. Antimicrobial susceptibility testing of Clostridioides difficile: a dual-site study of three different media and three therapeutic antimicrobials. Clin Microbiol Infect. 2025;31(6):1011–7 doi:10.1016/j.cmi.2025.01.028

24. Eucast. Clinical breakpoints 2025 [cited 2025 24-06-2025]. Available from: http://www.eucast.org/clinical_breakpoints/.

25. Vouga M, Greub G. Emerging bacterial pathogens: the past and beyond. Clin Microbiol Infect. 2016;22(1):12–21 doi:10.1016/j.cmi.2015.10.010

26. Advisory CTCa, (CTAG) G. Auxiliary Medicinal Products in Clinical Trials March 2024.

27. Boers SA, Hays JP, Jansen R. Novel micelle PCR-based method for accurate, sensitive and quantitative microbiota profiling. Sci Rep. 2017;7:45536 doi:10.1038/srep45536

28. Baloh M, Sorg JA. Clostridioides difficile spore germination: initiation to DPA release. Curr Opin Microbiol. 2022;65:101–7 doi:10.1016/j.mib.2021.11.001

29. Shen A. Clostridioides difficile Spore Formation and Germination: New Insights and Opportunities for Intervention. Annu Rev Microbiol. 2020;74:545–66 doi:10.1146/annurev-micro-011320-011321

